# Actin waves guide an outward movement of microclusters in the lymphocyte immunological synapse

**DOI:** 10.1101/2025.06.20.660656

**Authors:** Samuel Z Khiangte, Aheria Dey, Huw Colin-York, Marco Fritzsche, Sumantra Sarkar, Sudha Kumari

**Author notes:** co-first authors (Samuel Z Khiangte, Aheria Dey).

## Abstract

The lymphocyte immune response begins with antigen recognition on antigen-presenting cells, leading to the formation of the immunological synapse—a specialized interface for biochemical and biophysical exchange. At the synapse, most antigen-engaged receptor microclusters move inward toward the central supramolecular activation cluster (cSMAC) via retrograde F-actin flow, eventually clearing from the cell surface. This retrograde movement and receptor downregulation maintain antigen receptor homeostasis, critical for adaptive immunity, though its regulation remains unclear. Using live T cells, we identified a significant pool of antigen-engaged microclusters moving anterogradely toward the cell periphery, rather than the cSMAC. This movement was driven by actin waves propagating outward and coupling to microclusters through the Wiskott-Aldrich Syndrome Protein. These findings reveal a previously unrecognized mode of actin dynamics—anterograde actin waves—that co-exist with retrograde flow and direct microclusters away from the downregulation zone. This dual actin behavior underscores the complex cytoskeletal mechanisms T cells employ to regulate receptor distribution and maintain signaling homeostasis during immune activation.

## Introduction

The formation of a specialized cell-cell contact interface between the T cells and antigen-presenting cell (APC), the immunological synapse, signifies the initiation of adaptive immune response. Synapse allows the engagement of receptors and ligands between the cell surfaces, enabling molecular recognition events, including the engagement of T cell receptors with ligands on the APC, including major histocompatibility complex molecules loaded with agonist peptides. The engaged T cell receptors serve as signaling units and form microclusters that translocate within synapses ^1–3^. In its mature form, the synapse exhibits a bull’s eye organization with three distinct radially concentric zones ^4,5^ termed central supramolecular activation zone (cSMAC), peripheral supramolecular activation zone (pSMAC), and distal supramolecular activation zone (dSMAC) respectively. The T cell receptor engagement and activation occur in the dSMAC and pSMAC zones and the engaged receptors then migrate in a centripetal fashion traversing pSMAC to finally accumulate in cSMAC ^2,2^. At cSMAC, a major fraction of the antigen receptor exists in extracellular vesicles, with little or no signaling. The delivery of the antigen receptor to signaling poor cSMAC serves a crucial role in antigen receptor desensitization as well as extracellular communication^6–9^. Overall, the sub-synaptic positioning as well as dynamics of engaged receptors in SMACs determine some of the essential qualitative features of the T cell immune response ^2,10–14^.

Given its crucial importance in T cell receptor homeostasis at the synapse, the mechanisms of T cell receptor translocation from the periphery to cSMAC have been studied previously and found to be dependent on actin organization and dynamics^10,15,16^. Indeed, T cell receptor microclusters’ migration velocity correlates with the velocity of the centripetal flow of actin^15–19^, and breaking centripetal actin flow using micropatterned substrates leads to a disruption in centripetal microcluster trajectories and accumulation of T cell receptor at the actin flow breakpoints^15,18^.

Do all engaged T cell receptor microclusters undergo centripetal translocation and accumulate into cSMAC? While synapse imaging studies thus far indicate so, the surface expression analysis of antigen receptors using flow cytometry has shown an average of 40-60% of antigen-induced downregulation^20,21^ implying that on average ∼ 50% of antigen receptors escape the degradative fate. We investigated this inconsistency to examine if indeed the antigen receptor undergoes a ‘directional sorting’ at the immunological synapse during T cell activation. Using live primary T cells activated on APC-mimetic reconstituted bilayers, in combination with fast super-resolution imaging, unbiased tracking algorithm, and pharmacological inhibitors, we find that a sizeable fraction of motile clusters translocate not towards cSMAC but away from cSMAC. This motion is enabled by a previously uncharacterized wave-like anterograde flow of F-actin, which couples with a fraction of microclusters using WASP, sweeping them away from cSMAC towards the cell periphery. These results uncover a novel cytoskeletal behavior of T cells implicating the diversity of cell-intrinsic mechanisms mediating TCR homeostasis in the wake of an immune response.

## Results and Discussion

To generate planar T cell synapses that can be imaged at high spatiotemporal resolution for tracking ligated T cell receptor microcluster (referred to as “TCR” henceforth) movement, we utilized the supported lipid bilayer-based T cell activation system that has been previously used to examine subsynaptic receptor dynamics and movement ^15,16,22–26^. Mouse primary CD8+ T cells (see ‘Methods’) or the CD8+ 1G4 Jurkat T cell line ^27,28^ were incubated with supported lipid bilayers (SLB) reconstituted with a 6X-Histine tag containing the extracellular domain of ICAM1 and biotinylated and Alexa568 labeled anti-CD3 agonist antibodies (Okt3 and 2C11 clones respectively), and the SLB-T cell conjugates were imaged live using Total Internal Reflection – Structured Illumination (TIRF-SIM) microscopy (Figure 1A; ‘Methods’ section). The acquired images were processed (see ‘Methods’) and analyzed using dynamic tracking and particle image velocimetry algorithms (Python). The image processing scheme sensitively detected microclusters within the synapses with an accuracy of 97.2% (Figure 1B, Supp. fig. 1A). Detected microclusters were tracked using TrackPy to assess the magnitude and direction of all detected motile microcluster entities, and tracking accuracy was verified using independent manual tracking, where the velocities were found to be comparable (46.23 nm/s using Python and 49.18 nm/s using manual tracking) (Supp. fig. 1B, Movie 1). Analysis of TCR tracks showed a clear net centripetal movement (“retrograde or inward fraction”) in Jurkat T cells (Figure 1C-E, Movie 2), as described previously ^2,16,17,28–30^. However, when a similar analysis of TCR movement was performed in primary T cells, it revealed a significant deviation from the retrograde movement trajectories, where ∼40% of motile TCRs showed a net outward (“anterograde”) movement as well (Figure 1C-E, Movies 3 and 4), while overall TCR velocities were comparable in Primary and Jurkat T cells (Figure 1F). To examine if the outward fraction of TCR constitutes free anti-CD3 antibodies on the bilayer that are expected to exhibit multidimensional diffusion on the bilayer, we plotted the TCR directional fractions against intensities, given that free-anti-CD3 monomers are likely to be of lower intensities. We found that the directional fraction of TCR was independent of intensities (Figure 1G), indicating that TCR microclusters do contribute to both anterograde as well as retrograde fractions. TCR microclusters are known to form preferentially in pSMAC and dSMAC and then migrate towards cSMAC^2,5,10,31^. To examine if the anterograde TCR movement originates in a specific subsynaptic zone, we performed TCR movement analysis within radially segmented sub-synaptic zones. The results showed a lack of bias for outward movement in any subsynaptic zone, indicating that the anterograde movement of TCR can originate anywhere between cSMAC and the cell periphery (Figure 1H). Together, these data indicate that while the movement of engaged TCR is primarily centripetal in Jurkats, a significant fraction translocates outwards in primary T cells – a behavior that can originate anywhere across the synaptic interface.

**Figure 1:**
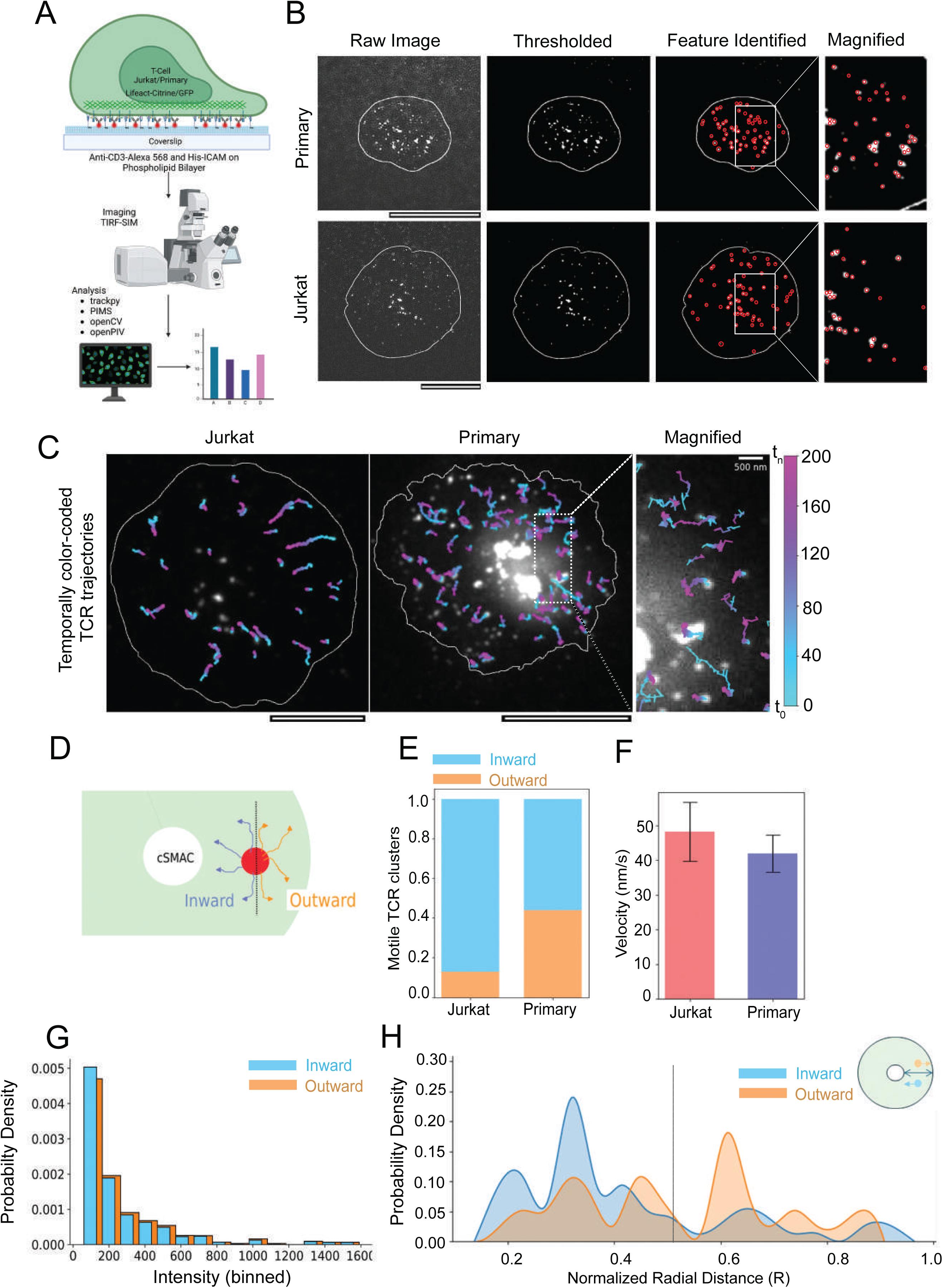
Live imaging of T cell immunological synapses using Total Internal Reflection-Structured Illumination Microscopy (TIRF-SIM) at superresolution scale to examine TCR microcluster movement dynamics. A. Schematic of the experimental setup utilized to visualize and analyze immunological synapses in live cells. The list shows the software employed in the analysis pipeline (see ‘Methods’). B. Examples of image processing steps underlying automated microcluster processing and detection in TIRF SIM images. C. pseudocolored images showing trajectories of TCR clusters tracked within 200 sec. The trajectories were color-coded based on their position in individual frames in reference to the initial frame. The reference color bar is shown on the right, where t_0_ refers to the time of initiation of imaging (t_0_ = synapse duration of <120 sec), and t_n_ refers to the time in the nth frame. D-F. A schematic showing the determination of “inward” (retrograde) and “outward” (anterograde) motion of TCR (D), and quantitative estimates of direction (E) and velocity (F) of individual TCR across 544 (Jurkat) and 1124 (Primary) TCR microclusters pooled from ≥12 cells in each case. The *P* values of comparison between Jurkat and Primary cell data in E and F are 0.0017 and 0.185, respectively, obtained using the Mann-Whitney non-parametric two-tailed test. G. The directional fraction of TCR is independent of microcluster intensities. The Y-axis represents the probability density of feature intensities (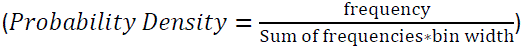), and the X-axis presents intensity bins. Note that the inward and outward fractions show a similar distribution of intensities (*p*-value of the comparison between the inward and outward fractions is 0.107, obtained using the Mann-Whitney two-tailed non-parametric test). H. Stratification of TCR tracks based on their position within radial zones of the synapse, where R=0 refers to the cSMAC boundary, and R=1 refers to the cell’s outermost boundary. The Y axis represents the probability density of TCR initial positions within the radial zones described in D; *p* value = 0.009 using Mann-Whitney non-parametric two-tailed test. Scale bar, 5µm.

The centripetal TCR flow is known to be guided by a retrograde F-actin flow, which arises via actin polymerization and rearrangement following antigen recognition^16,26,32^ ^35^. We wondered if the anterograde movement of TCR that we observed in primary T cells also relies on an outward actin flow, if any. First, to investigate whether the anterograde actin flows exist in the two T cell types, we imaged 1G4 CD8+ Jurkat T cell line stably expressing LifeAct-citrine^28^ or primary CD8+ T cells derived from lifeAct –GFP expressing mice^36^, since LifeAct has previously been used as a reliable marker of actin dynamics in immune cells^19,37–41^ (Figure 2A, Movies 5, 6). Indeed, we found the presence of an anterograde actin movement within the context of an overall retrograde flow at the synapse in primary T cells (Figure 2A, Movie 6). Both the velocity as well as the directions of the anterograde actin flow were comparable with the TCR outward fraction (Figure 2B). The actin anterograde flow was missing in Jurkats (Figure 2A, B, Movie 5). Thus, while the actin flow is primarily centripetal in Jurkats, directionally divergent actin flows exist in primary T cells with velocity and directions similar to that of TCR movement.

**Figure 2:**
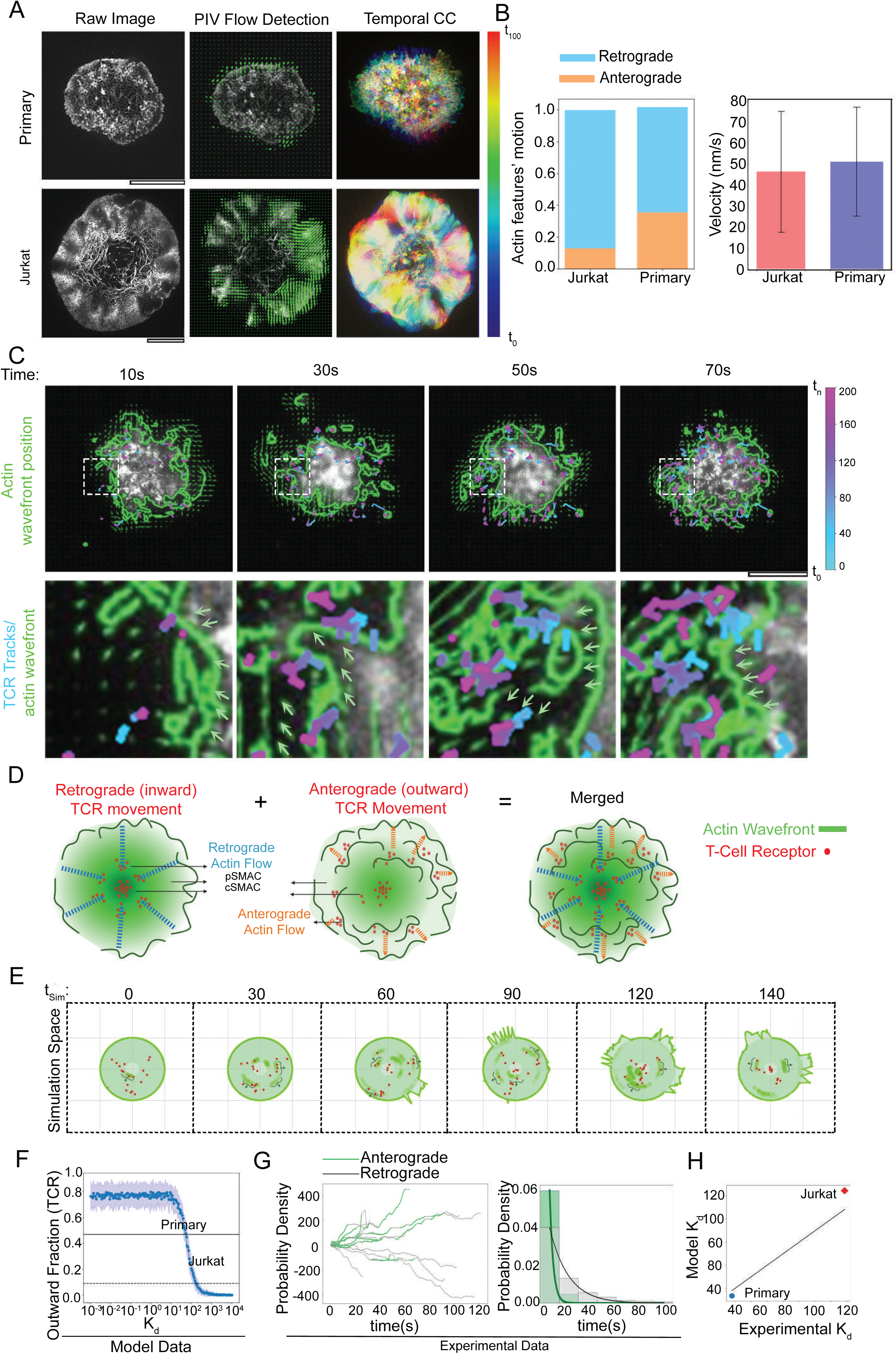
Actin dynamics at the synapse imaged using TIRF-SIM, and a computational model of actin-TCR interaction at the synapse. A. Particle image velocimetry (PIV) analysis of actin flows in primary (top middle panel) and Jurkat T cells (bottom middle panel). The arrows in PIV images show actin flow directions at a representative time point during the duration of imaging. The rightmost panels show temporal color coding of actin flows in the cells shown on the left. The color bar shows the pseudocolors corresponding to the relative temporal positions of actin features in the duration of 100 sec. Note that while the pseudocolored actin flows are smooth in Jurkat (bottom right panel), indicating a persistent motion, pseudocolored actin flows in the primary cell (top right panel) are heterogeneous, indicating microscale direction fluctuations. B. Quantification of directional fraction (left graph) and velocity (right graph) of actin features in Jurkat and Primary cells, (≥10 cells in each case; *p* values 0.0008 and 0.586 respectively, as obtained by Mann-Whitney non-parametric two-tailed test). C. Temporal sequence of actin dynamics at the synapse, with actin wavefronts’ positions are outlined with green lines (see ‘Methods’), and motile TCR marked as red dots. For clarity, the whole cell boundary is not shown in the images. D. A schematic showing the components of the computational model (actin waves, actin retrograde flow, and TCR at the synapse). The left panel highlights the retrograde movement of both actin and TCR, while the middle panel highlights their anterograde movement. The panel on the right shows both of the aforementioned components overlaid, as seen in primary T cells. E. Snapshots taken from the mechanistic computational model exploring the relationship between actin waves and TCR (red dots) to account for the behavior seen in C. The snapshots are taken from Movie 9. The relationship between TCR-anterograde actin binding affinity (K_d_) and overall TCR outward fraction, as predicted by the mechanistic model (F). G. Radial displacement values of individual TCRs, obtained from experimental data, where the initial position of all TCRs is taken as zero. A positive trajectory signifies anterograde TCR movement (green), whereas a negative trajectory (gray) signifies retrograde TCR movement. The plot represents TCR tracking values from a representative single cell. This analysis was performed for 12 cells, and a similar trend was seen for all cells. The plot on the right in (G) shows the probability density of the periods in inward (Gray and outward (Green) motion across all primary cells. H. A comparison of K_d_ values for primary and Jurkat cells estimated from the model with those obtained from the experimental time distributions shown in G (see methods). R^2^ for the fitted line in H is 0.883.

How could multidirectional flows, anterograde as well as retrograde, arise within the same subsynaptic locations? At the synapse, the retrograde actin flows are generated by actin polymerization at the plasmamembrane in the dSMAC zone followed by contraction of resultant F-actin by MyosinII^16,26,26,42^. To investigate the mechanism of anterograde flow generation at the primary T cell synapse, we closely examined the average behavior of overall actin flows at the synapses. We found distinct and repetitive “wavefront” –like structures arising and expanding outwards in the lamellar zone of the synapse. The actin wavefronts were associated with TCR outward fraction (Figure 2C, Movies 7, 8). Such wave-like excitations of actin cytoskeleton, although not superimposed on simultaneous and opposite flows, have previously been observed in other cellular systems, and are referred to as “actin waves” ^41,43–48^. These observations implied that the net TCR microcluster movement at synapse may rely on vectorially opposite flows from retrograde flow towards cSMACs and actin waves expanding away from cSMAC (Figure 2D). To gain further insights into this seemingly paradoxical observation, we explored the possibility of retrograde and anterograde flow co-existing and driving TCR microcluster movement within the synapse using a computational model. In this model, the anterograde flow is generated as a localized excitation within the cellular lamella at a random interval, and it propagates outward as a wavefront-like excitation with constant speed in randomly chosen directions (schematic in Figure 2E, Movie 9). When these wavefronts reach the cell periphery, they collide with and deform the cell boundary through elastic forces (Movie 9). The TCR are represented as featureless particles (red dots in Figure 2D) that can stochastically couple either to the retrograde flow (blue arrows in Figure 2D) or the anterograde flows (orange arrows in Figure 2D), although by default the TCR are coupled to the retrograde flow and migrate towards cSMAC. When bound to the anterograde flow, the clusters move away from the cell center; conversely, when they unbind from the anterograde flow, they instantaneously couple to the retrograde flow and start moving towards the cSMAC. The TCR can bind to the anterograde flow at a rate *k*_on_ and unbind from it with a rate *k*_off_. Since the exact values of *k*_on_ and *k*_off_ are not determined they both were treated as a free parameter. Assuming that the TCR-anterograde flow binding kinetics (order of msec) is much faster than the microcluster movement timescales (order of sec), we further presumed that the binding kinetics reaches chemical equilibrium within the observation timescale. In addition, since the binding affinity *K*_d_ (given as *K*_d_ = *k*_off_/*k*_on_*C*, where *C* is the concentration of F-actin in anterograde flow; *C* is taken as 100 μ*M*^49^)is not yet characterized, we treated it as a free parameter as well. Using these conditions, when we examined the changes in the TCR outward fraction, we found that it decreases monotonically with changes in *K*_d_ (Figure 2F).

Next, we compared the experimentally observed TCR outward fraction values with the ones in the simulated curve to estimate the binding affinity of TCR-anterograde flow (see ‘Methods’). The calculated TCR-anterograde flow binding dissociation constant (*K*_d_) values of primary cells (53.5 μM^−1^) were lower than the Jurkat cells (144.4 5 μM^−1^), implying that in addition to the presence of anterograde flow, the physical interaction of TCR with the anterograde flow may be required for the anterograde TCR movement in primary cells. To interrogate this hypothesis, we assessed the TCR-anterograde flow binding affinity from experimental data. For this, we analyzed the radial displacements of the TCR clusters in the experimental videos, and observed that when the TCR associated with the anterograde flows, it moved outward (green segments in the left graph in Figure 3G); otherwise, it moved inward (gray segments in the left graph in Figure 3F). To obtain the quantitative estimates of the binding affinity from TCR trajectories, we plotted the probability distribution of time the clusters spent in the anterograde and the retrograde directions (Figure 2G, graph on the right). The time spent in the anterograde flow-bound state was determined by the unbinding rate, k*_off_*, which we in turn estimated from the distribution of the green segments (Figure 3G left graph, Methods’). Repeating the analysis for the retrograde flow-bound (moving towards the c-SMAC) segments gave us an estimate for the binding rate, K*on* C. From these two estimates, we calculated the experimental dissociation constant (*K*_d_) for primary and Jurkat T cells to be 58.06 5 μM^-1^ and 128.2 μM^−1^, which matched well with the model estimate (53,5 μM^−1^ and 144.4 μM^−1^ (Figure 2H). This comparison evoked the hypothesis that even if the TCR has the potential to follow the anterograde flow, a coupling of the TCR to the anterograde flow is essential for its outward movement.

**Figure 3.**
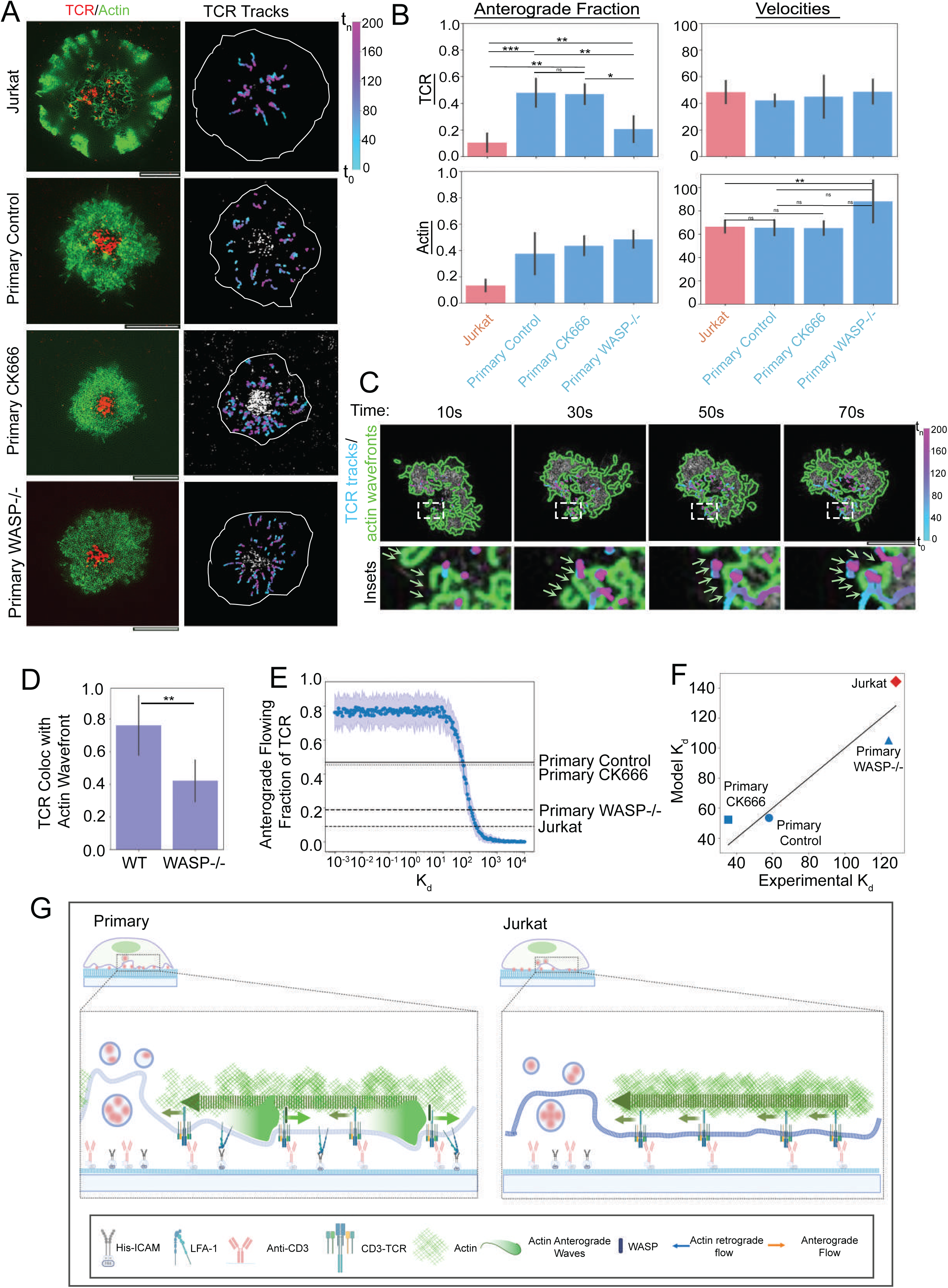
Molecular determinants of TCR-anterograde flow coupling at the immunological synapse. Imaging synapses of LifeAct expressing Jurkat (top panel) or primary T cells ICAM1 from the indicated backgrounds (bottom three panels) on SLBs containing anti-CD3-Alexa568 and using TIRF-SIM. The panels on the left show the final snapshot from the videos showing gross TCR and actin distribution, while the panels on the right show tracks of TCR clusters throughout the videos. B. Quantification of TCR motility (top two graphs) and actin flows (bottom graphs) showing directional fraction (left) and velocities (right). At least 10 cells were imaged in each case, and the p-values of the distribution are shown in Table 2 below, as obtained using the Mann-Whitney non-parametric two-tailed test. C. Tracking of actin waves (outlined in green lines) and TCR (marked as red dots) in LifeAct-GFP expressing WASP−/− primary T cell synapses. The wave-like behavior of actin was observed in WASP−/− T cells 80% of the time (total cells examined =15). D. Determination of TCR-wavefront colocalization in WT or WASP−/− primary T cells, as described in the ‘Methods’ section. For the analysis, a total of 10 cells were used, and *the p-*value of comparison is 0.0012, as determined using the Mann-Whitney two-tailed non-parametric test. E. An estimation of Anterograde TCR fraction using the computational model (see ‘Methods’). The fraction decreases monotonically with increasing *K*_d_ values. The experimentally observed fractions under various conditions are marked with straight lines, which shows that the anterograde fraction does not change with CK666 treatment, but does change for WASP−/− cells. F. A comparison of experimentally deduced and computationally derived *K*_d_ values across the specified perturbation backgrounds. The R^2^ value of the fitted line is 0.851. G. A schematic of TCR movement and underlying actin flows in primary T cells (left panel). Primary T cells exhibit retrograde actin flow to the cSMAC that drives TCR microcluster movement to the cSMAC via an unknown coupling mechanism. Overlaid on the retrograde actin flows, primary T cells exhibit anterograde flows composed of an ‘actin wave’ that expands towards the cell periphery and crashes on the cell boundary, resulting in lamellar extensions and retractions. Actin waves also sweep a sizeable fraction of TCR microclusters, with them leading to their anterograde movement away from cSMAC. The coupling of anterograde waves to TCR requires the actin nucleation promoting factor WASP, and lack of WASP while has no significant effect on actin waves, results in loss of TCR – actin waves coupling and subsequently, TCR anterograde movement. The Jurkat cells, unlike the primary T cells, exhibit dominantly the retrograde TCR and actin movements (panel on the right).

Table 2:

Table 2:
| Comparison | P-Value |
| --- | --- |
| **TCR Direction** |  |
| Jurkat WT vs. Mouse WASP-neg | 0.029154647 |
| Jurkat WT vs. Mouse Wt | 0.000179623 |
| Jurkat WT vs. Mouse ck666 | 0.000439567 |
| Mouse WASP-neg vs. Mouse Wt | 0.000442586 |
| Mouse WASP-neg vs. Mouse ck666 | 0.000922886 |
| Mouse Wt vs. Mouse ck666 | 0.928720781 |
| **TCR Velocity** |  |
| Jurkat WT vs. Mouse WASP-neg | 0.893918704 |
| Jurkat WT vs. Mouse Wt | 0.18571128 |
| Jurkat WT vs. Mouse ck666 | 0.449868171 |
| Mouse WASP-neg vs. Mouse Wt | 0.121989122 |
| Mouse WASP-neg vs. Mouse ck666 | 0.382284382 |
| Mouse Wt vs. Mouse ck666 | 0.761826409 |
| **ACTIN Direction** |  |
| Jurkat WT vs. Mouse ck666 | 0.000998029 |
| Jurkat WT vs. Mouse WASP-neg | 0.001015402 |
| Jurkat WT vs. Mouse Wt | 0.000800281 |
| Mouse WASP-neg vs. Mouse ck666 | 0.051332466 |
| Mouse Wt vs. Mouse ck666 | 0.223273048 |
| Mouse WASP-neg vs. Mouse Wt | 0.067936295 |
| **ACTIN Velocity** |  |
| Jurkat WT vs. Mouse ck666 | 0.202675046 |
| Jurkat WT vs. Mouse WASP-neg | 0.010241633 |
| Jurkat WT vs. Mouse Wt | 0.586661212 |
| Mouse WASP-neg vs. Mouse ck666 | 0.019445039 |
| Mouse Wt vs. Mouse ck666 | 0.89 |

To address the hypothesis that a coupling between TCR and actin anterograde flow is essential for the anterograde movement of TCR, we explored the molecular mechanisms mediating TCR-anterograde actin flow coupling. We first tested the role of nucleation factor Arp2/3 complex as a coupling mediator, since it can associate with TCR microclusters on one hand and with F-actin as a nucleation factor on the other hand, using pharmacological inhibitor CK666 that prevents Arp2/3 complex activation in T cells^19,38,50^. Treatment of cells with 50μM CK666 did not alter TCR outward fraction compared to the DMSO-treated control cells, and the overall TCR migration direction as well as velocity remained significantly different from the largely retrograde TCR movement in Jurkat cells (upper panels in Figure 3A, upper graphs in Figure 3B; Movies 10-12). Consistently, F-actin flow velocity and directional fraction also remained unchanged in CK666-treated cells (lower graphs in Figure 3B). These results indicated that branched actin nucleation via the Arp2/3 complex does not spontaneously guide the anterograde microclusters via anterograde flow. Next, we tested if the Wiskott-Aldrich Syndrome Protein (WASP) could enable the TCR-anterograde flow coupling since WASP can directly interact with TCR microcluster signalosome proteins ^51–53^, as well as with F-actin^54^. We examined TCR motility dynamics in WASP−/− primary CD8+ T cells expressing LifeAct-GFP. The WASP−/− cells showed significantly lower anterograde movement of TCR when compared to control cells (lower panels in Figure 3A, upper graphs in Figure 3B, Movie 13), and even though the anterograde flow and actin waves were still preserved in these cells (lower panels in Figure 3B, Supp. fig. 2), there was a lack of correlation between TCR movement and actin wavefronts (Figure 3 C, D; Movies 14, 15). These results indicated that WASP influences TCR anterograde movement without a direct role in generating the anterograde flow of actin. The simulations of TCR-actin flow interaction using our mechanistic model also corroborated the role of WASP as a coupler, where for a wide range of fixed parameters the Actin anterograde flow-TCR binding probability was around 0.05 for Jurkat cells and 0.5 for primary cells but reduced to 0.2 when the coupler was missing in primary cells (Figure 3E). These calculated values were comparable to the values obtained in WASP−/− cells (“WASP−/−” in Figure 3F; Supp. fig. 3). Thus, our data collectively suggests that WASP facilitates the coupling of TCRs to the anterograde actin flow.

Using an *in vitro* SLB-based ligand reconstitution system that has previously been extensively utilized to study synapse signaling, in combination with Primary T cells, our results reveal three novel aspects of antigen receptor and cytoskeletal behavior at the immunological synapse (Figure 3G): 1-there is an outward movement of TCR as well as F-actin at the synapse, 2-the generation of outward traveling actin waves that form simultaneously with the inward flow of actin, a phenomenon that could explain the simultaneous anterograde as well as retrograde microcluster movement, and 3-a key role of WASP in coupling receptor microclusters with actin waves for anterograde movement. The traveling actin waves have been described in a few cellular systems previously^44,46,55–57^, the unique feature in lymphocyte synapse is that here they exist simultaneously, in space and time, with a persistent retrograde actin flow. Whether the generation and sustenance of the two vectorially opposite actin flows at the same subcellular location follows cytoskeletal design principles unique to immune cells, and their underlying molecular ramifications that enable this highly counter-intuitive and specialized cytoskeletal dynamics, will be a subject of future research.

Surprisingly, in the Jurkat cell line, which has been a system of choice to investigate T cell activation, the majority of microclusters move only centripetally towards cSMAC. While the reason for the centripetal TCR migration bias is likely to be the absence of actin waves in Jurkat cells, why the actin waves fail to form in Jurkats is not clear at this stage. A lack of actin waves and TCR anterograde fraction in Jurkats-the most widely used cellular system for T cell biology-also explains at least to some extent why the anterograde microcluster movement has not been observed previously. In addition, the higher spatiotemporal resolution afforded by TIRF-SIM enabled the identification of fast microscale F-actin dynamics at early T cell synapse combined with unbiased cluster tracking, which could potentially be missed by routine cluster imaging and tracking methodologies. The actin waves described here can now be added to the microscale actin organizations at the immunological synapse that regulate the signaling architecture of T cells during activation^16,19,42,58^.

An interesting and somewhat confounding result we obtained was the selective use of WASP for coupling microclusters only to anterograde flow and not to the retrograde flow. The generation and sustenance of retrograde actin flow that guides the retrograde movement of clusters is known to be mediated by the formation of actin arcs and their contractions^16,17,26^. We found that while the retrograde flow is still the major contributor to the cluster migration, it seems to be coupled to the microcluster independently of WASP. The identity of the molecule/s that couple TCR to the actin arcs and retrograde flows in general, and the physical principles sustaining the coupling in both antegrade as well as retrograde movement will be a subject of future research.

Although we did not investigate the immunological significance of the anterograde microcluster movement directly, several elegant studies have demonstrated the significance of TCR movement in synaptic context ^6,9,52^ including its role in antigen receptor signal desensitization, degradation, exocytosis, and inter-cellular communication. The anterograde movement may alter or at least delay a sizeable fraction of TCR from these eventualities. Another tempting possibility is that the anterograde co-movement of microclusters along with actin waves prolongs their signaling lifetime, and may even augment signal amplification via long-range cytoskeletal consolidation^60,61^. Similarly, the sustained generation of actin waves and subsequent lamellar extensions may increase the cell surface area of synapse for better coverage of antigens on opposing interface for further TCR-MHCp engagement events ^62–66^. Finally, the propagation of lamellar waves will also likely impose mechanical forces on pre-existing as well as newly formed ligand-receptor bonds, including the TCR-MHCp pairs. Whether and how all of the above exciting possibilities combine at the synaptic interface to impact overall T cell antigen sensitivity, affinity discrimination, and signaling lifetime – essential features of early lymphocyte activation that rely on antigen receptor dynamics-will be a focus of future studies.

## Methods

### Cell isolation, culture, activation, and pharmacological treatments

Mouse CD8+ T cells were isolated either from wild type C57/BL6 mice, or from wild type C57/BL6 and WASP−/− C57BL6 mice expressing LifeAct-GFP, using the CD8+ T Cell Isolation Kit (Stem Cell Technologies) following the manufacturer’s protocol. The cells were cultured in sterile RPMI-1640 medium (GIBCO), supplemented with 10% fetal bovine serum (GIBCO), 2 mM L-glutamine (Thermo), 1 mM sodium pyruvate (Thermo), and 1% Penicillin/Streptomycin solution (Thermo). Post isolation, the cells were activated using PMA/Ionomycin (eBioscience™ Cell Stimulation Cocktail 500X), since activation using anti-CD3/CD28 is severely compromised in WASP-deficient CD8+ T cells^36^. The cells were maintained at a density of 0.5-2 x 10^6 cells/ml in a 37°C incubator with 5% CO_2_ and humidity. The Jurkat T cells utilized in this study were the 1G4 CD8+ T cell receptor variant, where the original endogenous T cell receptor was replaced human NY-ESO-sepcific CD8+ T cell receptor^27^. The 1G4 Jurkat cells were lentivirally transduced to generate stable cell lines expressing Lifeact-citrine^29^, and maintained in RPMI culture media mentioned above. At the time of imaging, the culture media was replaced with imaging media (phenol red –free *Ex vivo* 10 media (Lonza) supplemented with IL2 (10 units/ml), Fetal Bovine Serum (10%), and HEPES (10mM), and live imaging was performed in imaging media.

### Glass-supported lipid bilayers reconstituted with ligands

The supported lipid bilayers (SLBs) were prepared as described previously^23^. Briefly, the bilayers were deposited on 25mm round #1.5 glass coverslips (VWR), which were cleaned with piranha solution (sulfuric acid: hydrogen peroxide, 3:2 ratio) for 30 min, washed extensively with MilliQ water, and allowed to dry with ambient air at room temperature. The coverslips were then coated with a 5μl solution of equal volumes of DOPC liposomes (0.4 mM) and liposomes containing 12.5 mol% Ni2+ –NTA-DGS (0.4 mM), and 0.05 mol% cap biotin phosphatidylethanolamine (0.4 mM), and 37.5mol% DOPC (0.4 mM). The lipid film was then hydrated with HEPES Buffered Saline (20 mM HEPES, 140 mM NaCl, 5 mM KCl, 6 mM glucose, 1 mM CaCl2, 2 mM MgCl2; ‘HBS’) supplemented with 1% human serum albumin (HSA), pH 7.2., and washed several times with HBS. SLBs were then incubated with 5% bovine serum albumin supplemented with 100 μM NiCl_2_ and 1µg/ml streptavidin (to couple to biotin sites) for 30 min. SLBs were then washed extensively with HBS and incubated with monobiotinylated 2.5 µg/ml hamster anti-CD3ε (2C11 for murine cells and OKT3 for Jurkat cells) and 1 µg/ml 6X His-ICAM1 to activate T cells. The monobiotinylation of antibodies was assessed prior to the use using flow cytometric caliberation of ligand binding on glass bead supported lipid bilayers (Bangs Lab)^9,67^, and mobility of planar glass –supported bilayers was assessed prior to incubation with cells by examining the recovery profiles of Alexa568-anti-CD3 in photobleached zones on the lipid bilayers, and the recovery was found to be greater than 80% within 1 min of photobleaching in all experiments, consistent with high mobility of bilayers.

### Live-cell super-resolution extended TIRF-SIM

Extended total internal reflection fluorescence structured illumination microscopy (TIRF-SIM) was conducted using a 488-nm laser (Coherent, SAPPHIRE 488–500), which was directed through an acousto-optic tunable filter (AOTF, AA Quanta Tech, AOTFnC-400.650-TN). The laser beam was expanded and directed into a phase-only modulator composed of a polarization beam splitter, an achromatic half-wave plate (Bolder Vision Optik, BVO AHWP3), and a ferroelectric spatial light modulator (SLM; Forth Dimension Displays, SXGA-3DM). The diffraction pattern produced by the grating on the SLM passed through a polarization rotator, consisting of a liquid crystal variable retarder (LC, Meadowlark, SWIFT) and an achromatic quarter-wave plate (Bolder Vision Optik, BVO AQWP3). This configuration rotated the linear polarization of the diffracted light to preserve s-polarization, optimizing pattern contrast across all orientations.

To isolate the ±1 diffraction order, a hollow barrel mask driven by a galvanometer optical scanner (Model 623OH, Cambridge Technologies, Bedford, MA) was used to block higher-order diffraction light. The selected beams were then focused onto the back focal plane of a high-NA objective (Olympus Plan-Apochromat×100 Oil-HI 1.57NA) as two spots on opposite sides of the pupil. After collimation by the objective, the two beams interfered at the coverslip-sample interface at an angle exceeding the critical angle for total internal reflection, generating an evanescent standing wave of excitation that extended approximately 100 nm into the sample axially, with a laterally modulated sinusoidal pattern. This modulated pattern was a low-pass filtered, demagnified image of the grating displayed on the SLM. The emitted fluorescence was collected by the same objective, separated from the excitation light using a dichroic mirror, and imaged onto a sCMOS camera (Hamamatsu, Orca Flash 4.0 v2 sCMOS), where the structured fluorescence raw data were recorded. Cell samples were imaged in a micro-incubator (H301, okolab, Naples, Italy) under physiological conditions (37°C and 5% CO2). For each time point, three raw images were acquired at successive phase steps (0, 1/3, and 2/3) over the sinusoidal illumination pattern. This process was repeated with the excitation pattern rotated by +120° or −120° relative to the initial orientation. Phase stepping and pattern rotation were controlled by translating and rotating the grating image on the SLM. A total of nine raw images were acquired for each excitation wavelength before moving to the next. This acquisition cycle was repeated for each time point. Finally, the raw images were processed and reconstructed into SIM images. TIRF-SIM data were collected from at least 50 individual cells across three independent experiments.

### Data plotting and statistics

Data obtained using image analysis methodologies described in the aforementioned sections was plotted using either MATLAB or Prism (GraphPad). Cartoons and schematics were generated using BioRender. Figures were assembled using Adobe Photoshop (Adobe). Statistical comparison between groups was carried out primarily using Mann-Whitney two-tailed non-parametric distribution unless otherwise mentioned. The number of data points within distributions is mentioned in the figure legends.

### Tracking analysis

Images were analyzed using Python image processing packages trackpy^68^, openCV ^61^, and openPIV^69^. The videos are first processed using the Image J program: the images were converted to 8-bit grayscale image files, after which the images were filtered by applying a Gaussian filter of radius of 3 pixels (a window of radius 3 pixels was manually decided as it gave the best TCR detection when manually verified by colocalization with raw images to detect maximum clusters while avoiding artifacts (Supp. fig. 1A), when tested for 6 independent cells. The images were then thresholded to achieve individual microclusters as “particles” for tracking. To track TCR particle trajectories over time, we used dynamic tracking^68^. The probability for every new position of the particle was tracked using:

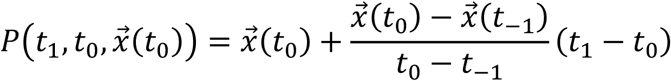

Where, 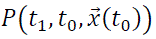 is the probability distribution of the particle at a time *t*_1_ given the particle was at 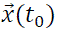 at time *t* = *t*_0_. Calculation of the probability requires calculating the instantaneous velocity, given by: 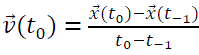. Here 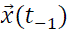 is the position of the cluster at a time previous to the current frame.

To ensure we track TCR microclusters and not any incidental free-floating debris on the SLB, we filtered out TCR trajectories with an end-to-end displacement greater than 3.5 pixels (at least 5% of the radius of the T Cell). We then examined the directionality of the detected TCRs. Finally, to ensure that the detected TCR features were in the dSMAC and pSMAC areas of active TCR transport, and not in cSMAC, where a large fraction of TCR accumulates, we utilized actin images as a positive mask for the tracked area, since actin cytoskeleton is depleted in the cSMAC zone^17,34^. To ensure we correctly tracked TCR clusters in the synapse, calculated trajectories were manually verified using ImageJ software (Supp. fig. 1B). Both methods gave comparable values for the velocity of TCR clusters which were further consistent with previously published values^29^. From the trajectory data, we could calculate the velocities and the directions of motile TCRs.

To quantify the fraction of inward and outward-moving TCR, we calculated the radial displacements of TCR from the cell center to the initial and final points in the trajectory. If the initial radial displacement was larger, the trajectory was classified as an ‘anterograde’ trajectory and vice versa. We also extracted spatial data for positional distributions of anterograde and retrograde TCR fractions shown in Figure 1H. To better visualize the distribution, we normalized the radial distances by the distance from the edge of the cSMAC to the cell periphery in all cells, such that the cell radius lay between 0 and 1 representing the periphery of the cSMAC and the cell periphery, in all cells. This allowed us to pool TCR trajectories across cells.

For tracking actin features in LifeAct images, the actin features were tracked to generate flow data. We used Particle Imaging Velocimetry (PIV) flow algorithms using the openPIV library in Python to quantify LifeAct flow directions and velocity. To validate the PIV tracking algorithm, we tested the results using two methods. First, we checked whether the algorithm gave retrograde actin velocities consistent with previous studies^29^. Second, we performed particle-based tracking of actin instead of flow-based analysis, as done for TCR clusters mentioned above. The actin images were processed using a Gaussian filter as described for TCR images, and again thresholded to pick out the brightest actin features. The directional fraction values from this method resulted in an average outward fraction of 0.44 as compared to 0.38 as done by PIV tracking.

Edge detection of actin wavefronts was performed using a Sobel edge detection algorithm^70^ to detect the actin-rich wavefronts. First, the image was processed through a 5×5 Gaussian filter^19^, which blurs the image so that only the most prominent edges are detected. Only potential edges within a predetermined minimum and maximum values of intensity gradients were filtered (see below). To determine the appropriate edge gradients we manually examined the pixel values of edge gradients of the wave features and took the average of intensities from 4 cells in each primary control and primary WASP−/− cells. From this measurement, we found the minimum edge gradient as 10 (arbitrary units, AU) and the maximum as 40 AU. We then assessed the colocalization of TCR with the detected actin wavfront features. If an edge coordinate (green points in Figure 2C) is present within eight nearest pixel neighbors of the TCR (red point in Figure 2C), the TCR was treated as colocalized with actin wavefront for that time point. This analysis was performed for entire image stream in a video, and the values of all TCR for all time points were averaged and presented in the graph (Figure 3D).

### Computational Modelling

We developed a phenomenological model of TCR transport via the actin cytoskeleton. It has already been shown in the Jurkat cells that TCR couples to the actin retrograde flow and moves persistently toward the cSMAC region of the immunological synapse. In the mouse primary cells, we found that the TCR movement correlates with both retrograde flow and anterograde flow. To model this system, we assumed that the TCRs are featureless particles that are confined to move on the plane of the synapse. In this model, the TCRs are constitutively coupled to the retrograde flow, so that they naturally move towards the cell center. In the course of their retrograde movement, the TCR can stochastically bind to (and unbind from) the anterograde flow as well, with a rate *k*_on_(*k*_off_). Following experimental observations, we modeled the anterograde flow as traveling wave-like excitations of the actin cytoskeleton. As a result, in the anterograde flow, the wavefront-like regions, rich in F-actin, propagate away from the cSMAC. In the model, the wavefronts are generated at a fixed rate from the cell center, while their propagation direction is uniformly and randomly chosen between 0 and 2π.

While bound to the anterograde flow, the TCR moves towards the direction of propagation of the wavefronts. Once unbound from anterograde actin flow, the clusters immediately rebind to the retrograde flow and start moving towards the cSMAC. Since the fraction of anterograde flow-bound trajectories would depend on the binding and unbinding rates, we characterized this dependence by measuring the fraction of anterograde flow-bound trajectories as a function of 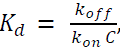, where *C* is the average concentration of F-actin in the cell (taken to be 100 μM, see **Table 1**), which showed a nonlinear dependence. For each value of *K*_d_, we compute the average fraction of anterograde flow-bound trajectories from 50 model trajectories. The other parameters are listed in Table 1 (Below).

|  |  |
| --- | --- |
| The radius of the cell (primary) | 5$\mu\text{m}$ |
| Inward Velocity of TCR | 0.075$\mu\text{m/s}$ |
| Number of TCR | 50 |
| Outward Velocity of waves | 0.1$\mu\text{m/s}$ |
| Frequency of wave generation | 0.4 /s (avg) |
| Concentration actin in a wave | 100$\mu\text{M}$ |

In the mechanistic model, the loss of coupling corresponds to the lowering of the binding probability (i.e., an increase in *Kd*, when all other parameters are fixed). To understand the dependence of outward fraction on Kd, we ran an ensemble of simulations for a wide range of *Kd* values, holding all other parameters fixed (initiation rate of the anterograde flow excitations and the average direction of its movement). We found that below a threshold *Kd* value (equal to the average actin concentration, which we take to be 100 μM^49^, the outward fraction stays at a high value first, and then above a threshold, it rapidly decays. Interestingly, the experimentally observed values of outward fraction were predicted at low *Kd* values for untreated and CK666-treated primary cells, whereas the Kd value is almost two-fold higher for WASP−/− cells. While these numbers were model-dependent, they did provide an estimate of the transport mechanism of the anterograde flow-bound TCR.

To validate the model, we plotted the persistence time distribution from the experimental data (Figure 2G, Supp. fig. 3) i.e. the distribution of the durations when the actin waves are pushing out the TCR clusters. Theoretically, this data is expected to follow an exponential distribution as shown in Equation 1. This allowed us to get an experimental estimate of *K*_off_. Doing a similar analysis for the switching time distribution, i.e., the distribution of the time the cluster is not being pushed out, allowed us to estimate *K*_on_ using equation 2, where C is the actin concentration.

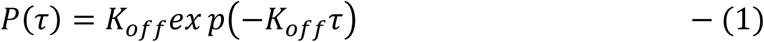

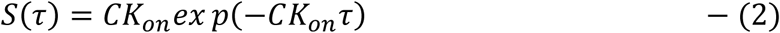

We then calculated 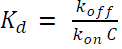 and compared the experimental rates with those predicted by the model. The comparison shown in Figures 2H and 3F indicates good agreement between *K_d_* values obtained experimentally with those derived using the model.

## Acknowledgements

We thank the Biology Divisional Microscopy facility and Central animal facility at the Indian Institute of Science; the Advanced Imaging Center at Janelia Campus, and J. Haddleston for help with the lattice light-sheet microscopy (The Advanced Imaging Center is jointly funded by the Howard Hughes Medical Institute and the Gordon and Betty Moore Foundation). SS acknowledges generous support from Axis Bank Center for Maths and Computing (OD/ACMC-23-0013) and SERB-DST India (SRG/2022/000163). Both SS and SK acknowledge generous support from the MoE-STARS grant. SK acknowledges ANRF/SERB grant (SPG/2021/004030), an Infosys Young Investigator fellowship, and an Intermediate Fellowship from India Alliance DBT-Wellcome Trust (IA/I/23/1/506757).

## Supplemental data

**Supplementary figure 1.** Validation of automated TCR microcluster (“feature”) detection (A) and tracking (B) routines, by either overlaying detected clusters with either raw images (A), or by comparing the TCR tracks obtained from manual and automated tracking (B). Manual tracking was performed in ImageJ, where a particle trajectory was plotted by manually identifying the position of the particle at each time and then integrating individual positions to generate the tracks.

**Supplementary figure 2** Probability densities of time spent in the anterograde (green) and retrograde (gray) direction by TCR clusters. These probability densities were obtained from Jurkat, WASP−/−, and CK666-treated cells, and are analogous to Figure 3F, which were made using data from WT mouse primary cells. The *p values* of comparison between groups were: Jurkat vs Primary WT 0.00216; Jurkat vs Primary CK666 0.00116; Jurkat vs Primary WASP−/− 0.00432; Primary WT vs Primary CK666 0.08158; Primary WT vs Primary WASP−/− 0.08650; Primary CK666 vs Primary WASP−/− 0.26767, as obtained using Mann-Whitney non-parametric two-tailed test.

**Supplementary figure 3**. (A) Kymographs showing actin waves in Jurkat (top panel) and primary T cells (bottom three panels), the green lines indicate the evolution of actin boundary over time. (B) A comparison of the fluctuation from the mean of the cell edges (RMS fluctuation of the green line; p-values are given in Table 2).

**Supplemental Movie 1.** Comparison of manual (left panel) and automated (Right panel) tracking of TCR clusters in Jurkat T cells.

**Supplemental Movie 2.** Automated tracking of TCR clusters in Jurkat T cells. The left panel shows raw images, while the panel on the right shows positional color-coded trajectories. The movie corresponds to Figure 1C.

**Supplemental Movie 3.** Automated tracking of TCR clusters in primary T cells. The left panel shows raw images, while the panel on the right shows positional color-coded trajectories. The movie corresponds to Figure 1C.

**Supplemental Movie 4.** Inset showing magnified details of TCR trajectories from Movie 3.

**Supplemental Movie 5.** PIV flow analysis of LifeAct-GFP in Jurkat T cells. The left panel shows LifeAct-GFP distribution, while the panel of the right panel shows corresponding PIV flow vectors (pseudocolored green) overlaid on LifeAct-GFP (greyscale) at the given time points.

**Supplemental Movie 6.** PIV flow analysis of LifeAct-GFP in primary T cells. The left panel shows LifeAct-GFP distribution, while the panel of the right panel shows corresponding PIV flow vectors (pseudocolored green) overlaid on LifeAct-GFP (greyscale) at the given time points.

**Supplemental Movie 7.** Actin wavefront detection and corresponding TCR microcluster trajectories in a primary T cell. The left panel shows TCR (pseudocolored red) and LifeAct (pseudocolored green) distribution in raw images, while the panel on the right shows an overlay of detected actin wavefrons (green lines) and TCR trajectories (positional color-coded tracks).

**Supplemental Movie 8.** Wavefront movement(green lines), corresponding PIV actin flow vectors (green arrows), and positional color-coded TCR microcluster trajectories in a primary T cell.

**Supplemental Movie 9.** A movie of the mechanistic model to explore actin waves – TCR trajectories. Although the model includes retrograde flow of actin, it has not been represented in the visual depiction of the simulation space. For details, see ‘Methods’.

**Supplemental Movie 10.** LifeAct-GFP (top left panel, pseudocolored green) and TCR (top right panel, pseudocolored red) movement in Jurkat T cell. The bottom left panel shows a combined distribution of the top two panels, while the bottom right panel shows positionally color-coded TCR trajectories obtained using automated tracking. The movie corresponds to Figure 3A.

**Supplemental Movie 11.** LifeAct-GFP (top left panel, pseudocolored green) and TCR (top right panel, pseudocolored red) movement in a DMSO-treated primary T cell. The bottom left panel shows a combined distribution of the top two panels, while the bottom right panel shows positionally color-coded TCR trajectories obtained using automated tracking. The movie corresponds to Figure 3A.

**Supplemental Movie 12.** LifeAct-GFP (top left panel, pseudocolored green) and TCR (top right panel, pseudocolored red) movement in a CK666-treated primary T cell. The bottom left panel shows a combined distribution of the top two panels, while the bottom right panel shows positionally color-coded TCR trajectories obtained using automated tracking. The movie corresponds to Figure 3A.

**Supplemental Movie 13.** LifeAct-GFP (top left panel, pseudocolored green) and TCR (top right panel, pseudocolored red) movement in a WASP-deficient primary T cell. The bottom left panel shows a combined distribution of the top two panels, while the bottom right panel shows positionally color-coded TCR trajectories obtained using automated tracking. The movie corresponds to Figure 3A.

**Supplemental Movie 14.** Actin wavefront movement (marked by the green line in the right panel) and TCR trajectories (positionally coded trajectories derived using automated tracking; right panel) in a WASP−/− primary T cell. The panel on the left shows the combined distribution of LifeAct (pseudocolored green) and TCR (pseudocolored red).

**Supplemental Movie 15.** Another example of actin wavefront movement (marked by the green line in the right panel) and TCR trajectories (positionally coded trajectories derived using automated tracking; right panel) in a WASP−/− primary T cell. The panel on the left shows the combined distribution of LifeAct (pseudocolored green) and TCR (pseudocolored red).

