## Supplementary figures and images for "Actin waves guide an outward movement of microclusters in the lymphocyte immunological synapse"

### Supplemental figures

#### Supplementary figure 1

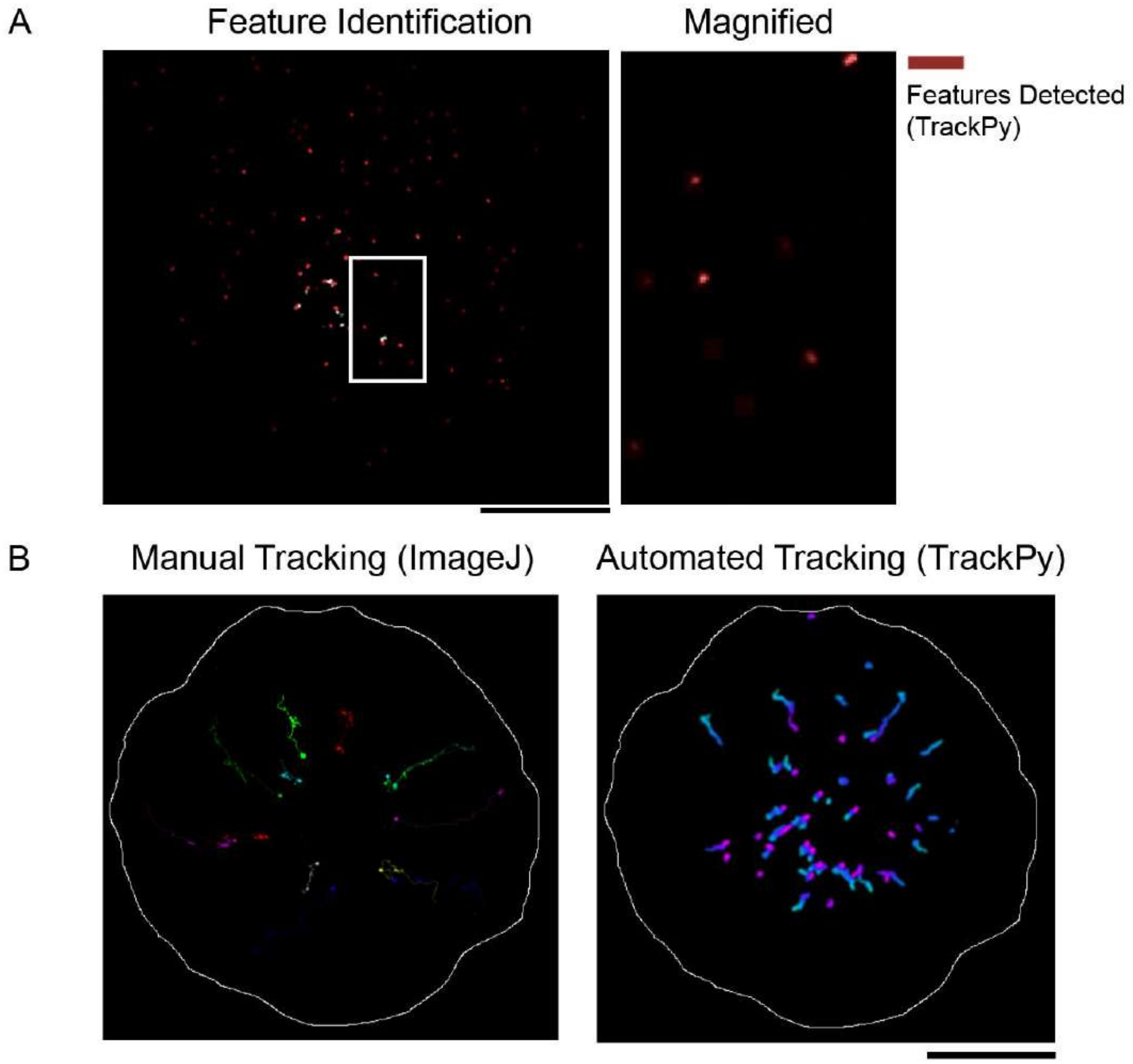

Supplementary figure 2

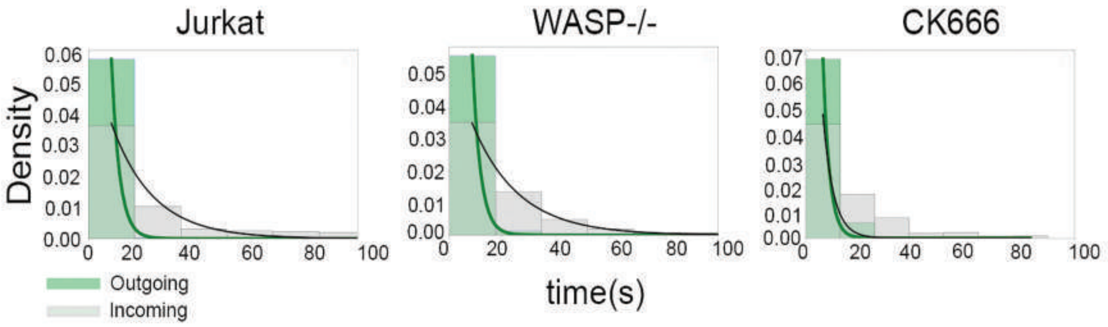

#### Supplementary figure 3

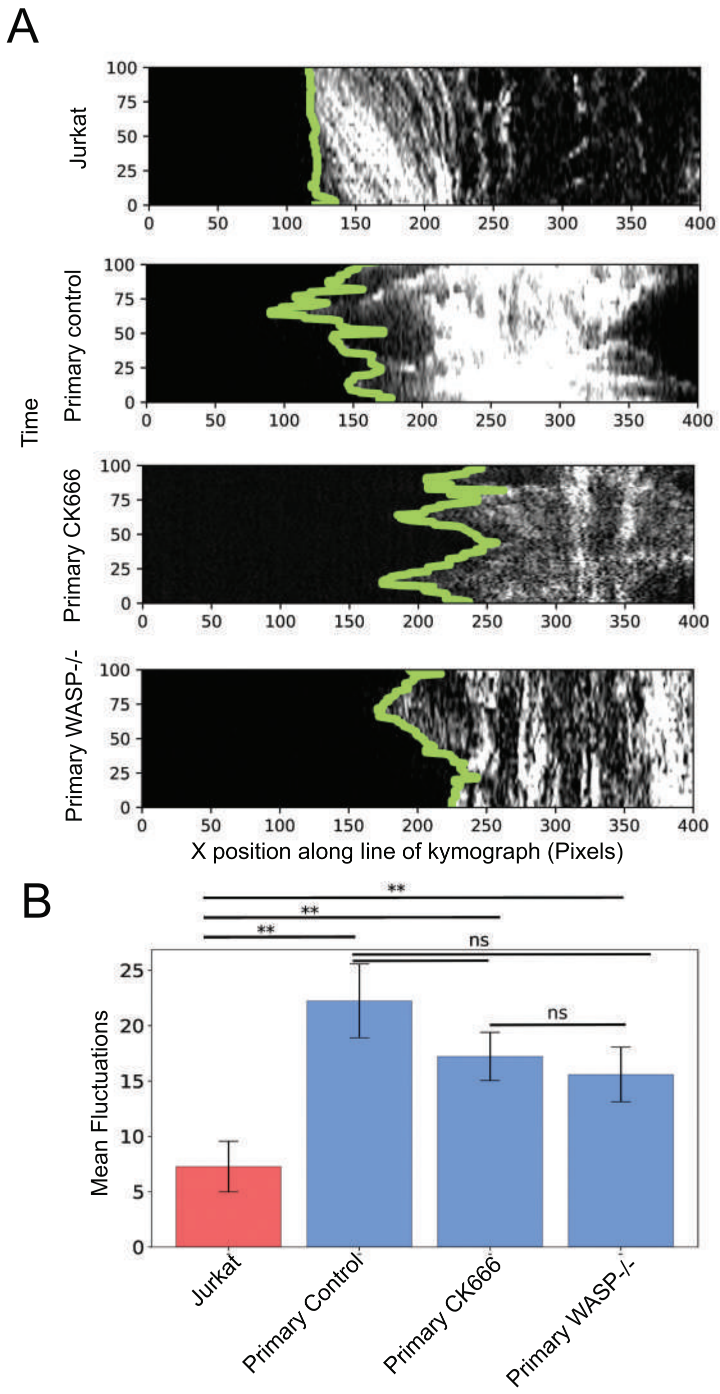
